# A high-quality reference genome for the common creek chub, *Semotilus atromaculatus*

**DOI:** 10.1101/2023.07.14.549000

**Authors:** Amanda V. Meuser, Amy R. Pitura, Elizabeth G. Mandeville

**Author notes:** Corresponding author*: Amy Pitura, University of Guelph, 50 Stone Road East, Guelph, Ontario N1G 2W1, Canada, 1-519-824-4120 ext. 52843. Co-first authors (Amanda V. Meuser, Amy R. Pitura).

## Abstract

Creek chub (*Semotilus atromaculatus*) are a leuciscid minnow species commonly found in anthropogenically disturbed environments, making them an excellent model organism to study human impacts on aquatic systems. Genomic resources for creek chub and other leuciscid species are currently limited. However, advancements in DNA sequencing now allow us to create genomic resources at a historically low cost. Here, we present a high quality 239 contig reference genome for the common creek chub, created with PacBio HiFi sequencing. We compared the assembly quality of two pipelines: Pacific Biosciences’ Improved Phase Assembly (IPA; 873 contigs) and Hifiasm (239 contigs). Quality and completeness of this genome is comparable to the zebrafish (Danioninae) and fathead minnow (Leuciscidae) genomes. The creek chub genome is highly syntenic to the zebrafish and fathead minnow genomes, and while our assembly does not resolve into the expected 25 chromosomes, synteny with zebrafish suggests that each creek chub chromosome is likely represented by 1-4 large contigs in our assembly. This reference genome is a valuable resource that will enhance genomic biodiversity studies of creek chub and other non-model leuciscid species common to disturbed environments.

## Introduction

Genomic studies of non-model species have become increasingly feasible in the past two decades (Narum *et al*. 2013, Lou *et al*. 2021). Efforts are now under way to sequence the tree of life (Fan *et al*. 2020), including organisms of no known economic importance, whose genomes are likely to be used primarily for conservation or evolutionary research. While genomic studies of non-model organisms are proceeding at a rapid pace, progress is still limited by the lack of suitable reference genomes for many species and clades. Using a reference genome from a closely related species is sometimes possible when there is no available reference for focal taxa (e.g. Mandeville *et al*. 2019). However, this approach can produce misleading results under some circumstances, including when the goal of a study is to examine within-species differentiation or similar and species-specific variation may be lost (as when wolf vs. dog reference genomes were used for a study of wolves; Gopalakrishnan *et al*. 2017). These concerns are especially relevant when considering clades where a substantial amount of structural genomic variation exists.

Fish genomes are quite diverse, and vary in size from 0.5–2 pg (excluding polyploids; Smith & Gregory 2009). On a locus-specific functional level, essential processes like sex determination can have an incredibly diverse genetic basis in fish (Bachtrog *et al*. 2014, Pennell *et al*. 2018). Within North American teleost fish, a large proportion of fish biodiversity is encompassed by the family Leuciscidae within the order Cypriniformes (Holm *et al*. 2022, Stout *et al*. 2016), but previous genomic work on leuciscid minnow species has been limited. However, being extremely numerous and geographically widespread, these species have great potential for use as model species to study the effects of anthropogenic disturbance and overall population genetic structure of stream fishes. Some species are quite tolerant, and persist or even thrive in disturbed environments (Stammler *et al*. 2008). Additionally, these species are known to hybridize from morphological data, but hybridization patterns have not been described in detail using genetic or genomic data (Corush *et al*. 2021).

One major limitation for future work is that no leuciscid reference genomes have been available until recently, and the highest quality available reference genome would be a zebrafish (*Danio rerio* genome), which is not very closely related to many wild leuciscid taxa of interest (Fig. 1; Schönhuth *et al*. (2018)). A recently published fathead minnow (*Pimephales promelas*) genome provides one resource in the leuciscid family (Martinson *et al*. 2022), but there is still a dearth of genomic resources for this hyper-diverse and geographically ubiquitous clade. In consequence, previous genomic studies of creek chub have relied on artificial reference genomes or reference genomes from distantly related taxa (e.g. Meuser *et al*. 2022).

**Figure 1.**
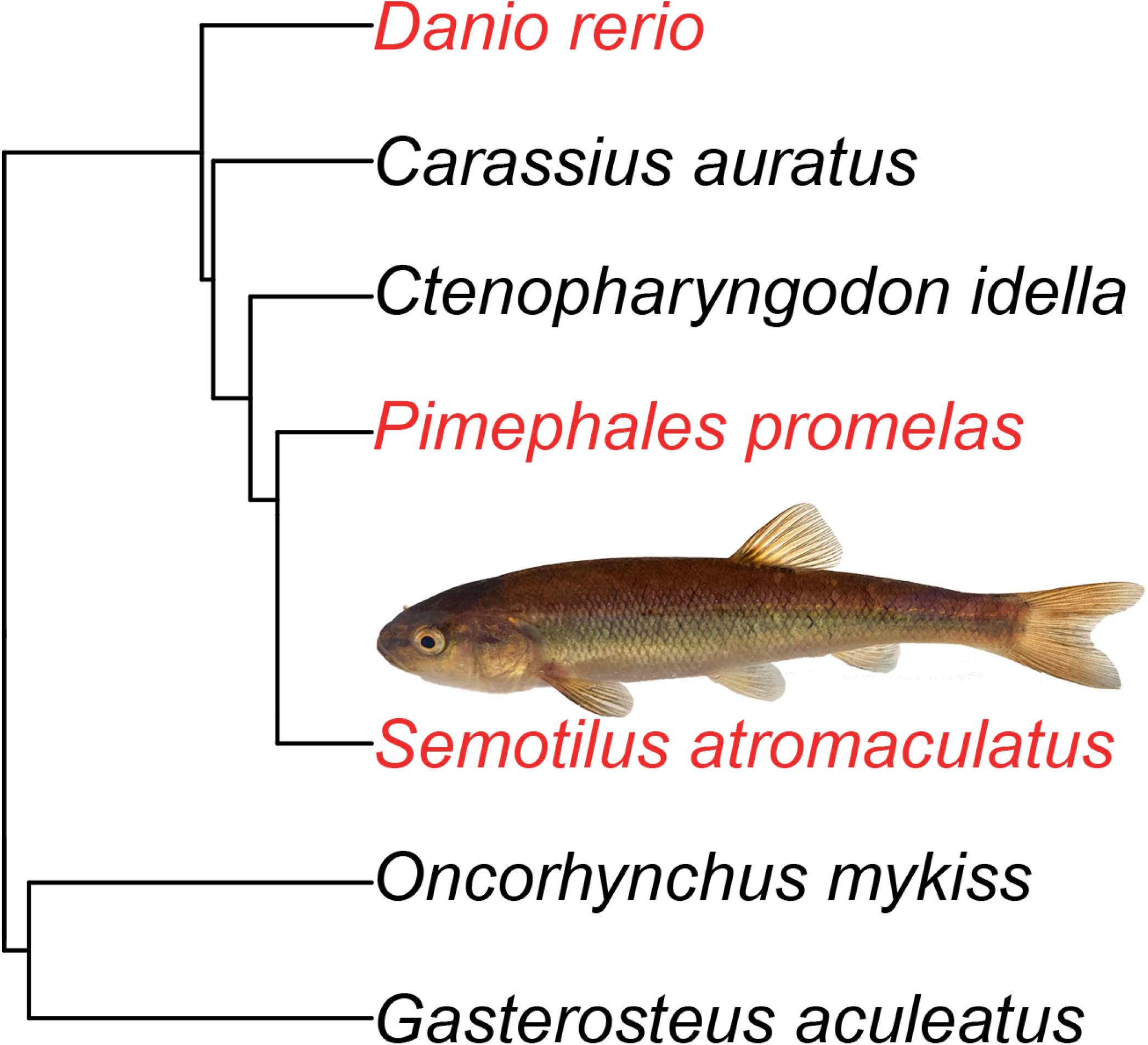
: Phylogeny showing the relationship between zebrafish, fathead minnow, creek chub, and other fish commonly used as model species. The inset photo shows a creek chub individual. The phylogeny was created using data from the fishtree package in RStudio (Chang *et al*. 2019).

We sequenced the genome of the common creek chub, *Semotilus atromaculatus*, to provide a genomic resource for future studies of creek chub and other leuciscid minnows in North America. We chose to sequence creek chub because of the broad range, abundant population sizes, and generally tolerant life history of this species. Creek chub are expected to have 2n=50-52 chromosomes and a genome size of 1.25pg, similar to many leuciscid species (Gold & Amemiya 1987, Legendre & Steven 1969). We also compared the outcomes of two assembly pipelines: Pacific Biosciences’ Improved Phase Assembly (IPA) HiFi Genome Assembler pipeline (github.com/PacificBiosciences/pbipa) and the software HiFiasm (Cheng *et al*. 2021). Finally, we assessed synteny between the creek chub reference genome and the zebrafish and fathead minnow genomes (Martinson *et al*. 2022).

## Methods

Sampling was accomplished under animal utilization protocol #4237 approved by the University of Guelph Animal Care Committee, with permits from the Ontario Ministry of Natural Resources and Forestry (Licence No. 1100698), and with private landowner permission. One wild-caught, 10.6 cm (total length) creek chub was sampled to generate this reference genome (see Fig S1 for a photo). We sampled this fish using a beach seine from Swan Creek in southern Ontario, Canada, in August 2022. The sampled individual was identified morphologically by expert field personnel and later identified genetically as a creek chub with DNA barcoding. Following capture, we euthanized the target individual with an overdose of MS-222, then sampled and flash froze muscle tissue in liquid nitrogen within 5 minutes of euthanasia to ensure preservation of high molecular weight DNA. We stored the flash-frozen muscle and remainder of the specimen in a -80°C freezer, except for 2 fin clips preserved in 95% ethanol. We extracted DNA from the fin clips using a DNeasy Blood & Tissue kit (Qiagen) and quantified the concentration using a NanoDrop 8000 Spectrophotometer (Thermo Scientific). We used this DNA to verify our phenotypic identification with DNA barcoding at the University of Guelph’s Advanced Analysis Center. The COI-3 region of the mitochondrial genome was amplified and sequenced using thermalcycler conditions and primers from Ivanova *et al*. (2007). The forward fasta sequence was input on BOLD’s Identification Engine (www.barcodinglife.org/index.php/IDS_OpenIdEngine; (Ratnasingham & Hebert 2007)) and confirmed to belong to creek chub.

We sent flash frozen muscle tissue to the University of Delaware’s DNA Sequencing & Genotyping Center, in Newark, Delaware, USA. High molecular weight (HMW) DNA extraction was completed using the MagAttract HMW DNA kit (Qiagen), then the extracted DNA was quantified using a Qubit Fluorimeter and DNA fragment sizes were assessed by Femto Pulse system instrument (Agilent). Next, a Megaruptor 2 (Diagenode) was used to shear 3 *μ*g of DNA to 15kb fragments. Then, a SMRTbell DNA library was constructed according to the Pacbio HiFi SMRTbell protocol using SMRTbell Express Template Prep Kit 3.0 (Pacbio, 102-182-700). After BluePippin size selection (Sage Science, PAC20KB) removed fragments smaller than 8 kb, the average size in the library was 18 kb based on Femto Pulse System (Agilent) analysis. Finally, sequencing was performed on 2 SMRT 8m cells on Sequel IIe instrument with 30 hours movie, using both the Sequel II Binding kit 2.2 and Sequel II Sequencing kit 2.0.

The initial assembly of the reference genome was performed by the University of Delaware’s DNA Sequencing & Genotyping Center. They used Pacific Biosciences’ Improved Phase Assembly (IPA) HiFi Genome Assembler pipeline (github.com/PacificBiosciences/pbipa). In addition, we used HiFiasm (Cheng *et al*. 2021) to create a genome assembly. This, and all subsequent computation, was performed on Digital Research Alliance of Canada’s Cedar high performance computing cluster. We ran HiFiasm (v0.16.1) with 32 CPUs to create a HiFi-only assembly, as we did not have parental short reads or Hi-C reads to create either the trio-binning or Hi-C integrated assemblies (Cheng *et al*. 2021).

We assessed genome assembly quality using custom R and shell scripts to quantify distribution of assembled contig and scaffold lengths, and the number of unique assembled scaffolds (Fig. 2). From these data, we calculated N50, N90, L50, and L90, as well as maximum, mean, and median contig length (Table 1). As this assembly is comprised completely of long-read PacBio data, there are no gaps in our assembled contigs, and we hereafter refer to these fragments of the genome simply as contigs. We also ran this analysis on the most recent versions of the zebrafish (GCF 000002035.4 GRCz11) and fathead minnow (GCF 016745375, Martinson *et al*. 2022). We assessed completeness of the creek chub reference genome using BUSCO v5.2.2 with actinopterygii odb10 as the database (Sim ã o *et al*. 2015), as well as the zebrafish and fathead minnow reference genomes for the sole purpose of comparison with the same database. We used kraken 2 v2.1.2 (Wood *et al*. 2019) to assess contamination using a custom database containing virus, plasmid, protozoa, archaea, bacteria, human, plant, and fungi sequences.

**Table 1:**
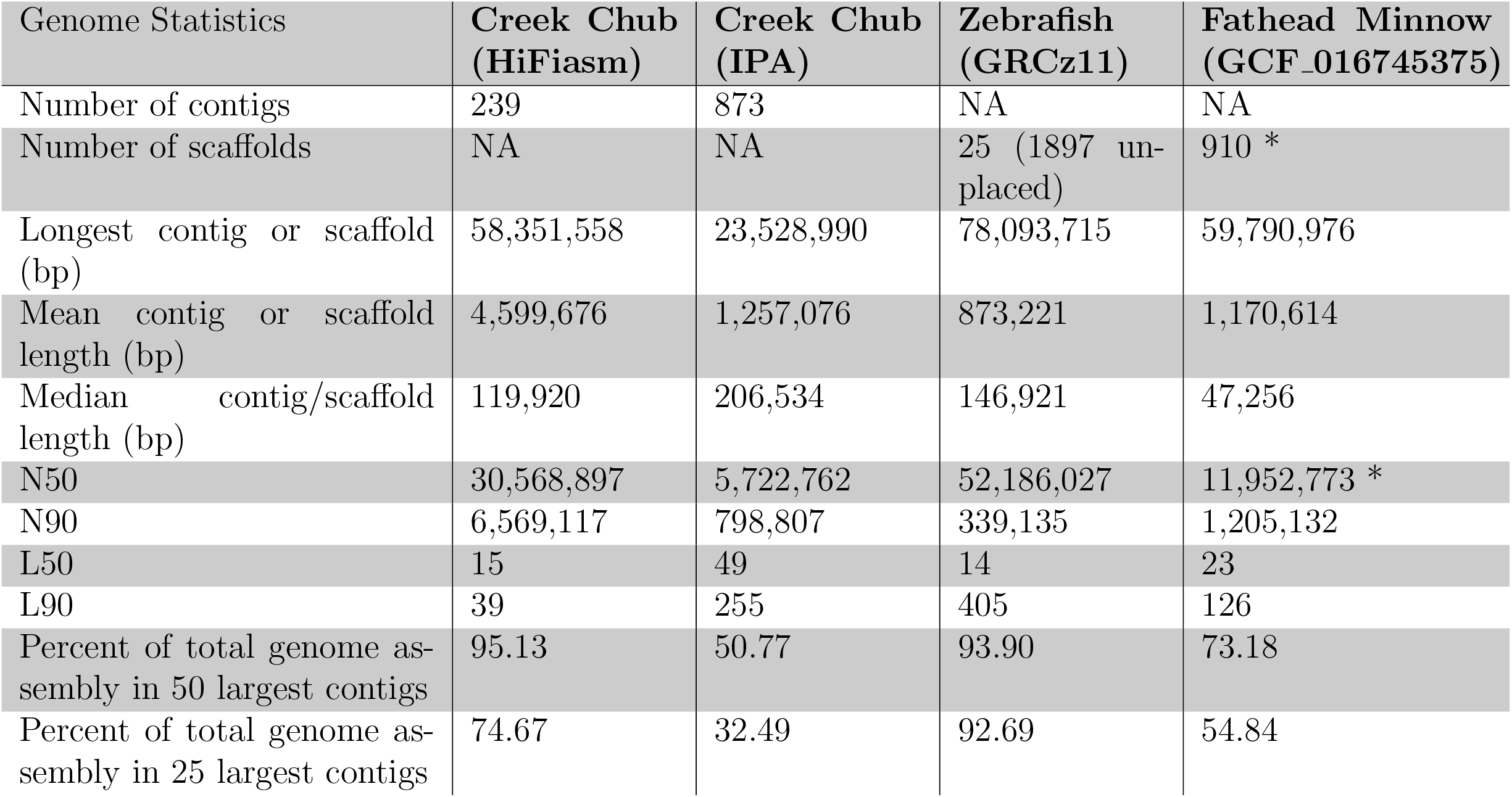
Genome statistics for both the HiFiasm and IPA genome assemblies and the most recent versions of the zebrafish and fathead minnow genomes. Statistics for each assembly were generated using a custom script written in a combination of both shell and R. NA = not applicable. * Denotes fathead minnow statistics from Martinson *et al*. (2022).

**Figure 2.**
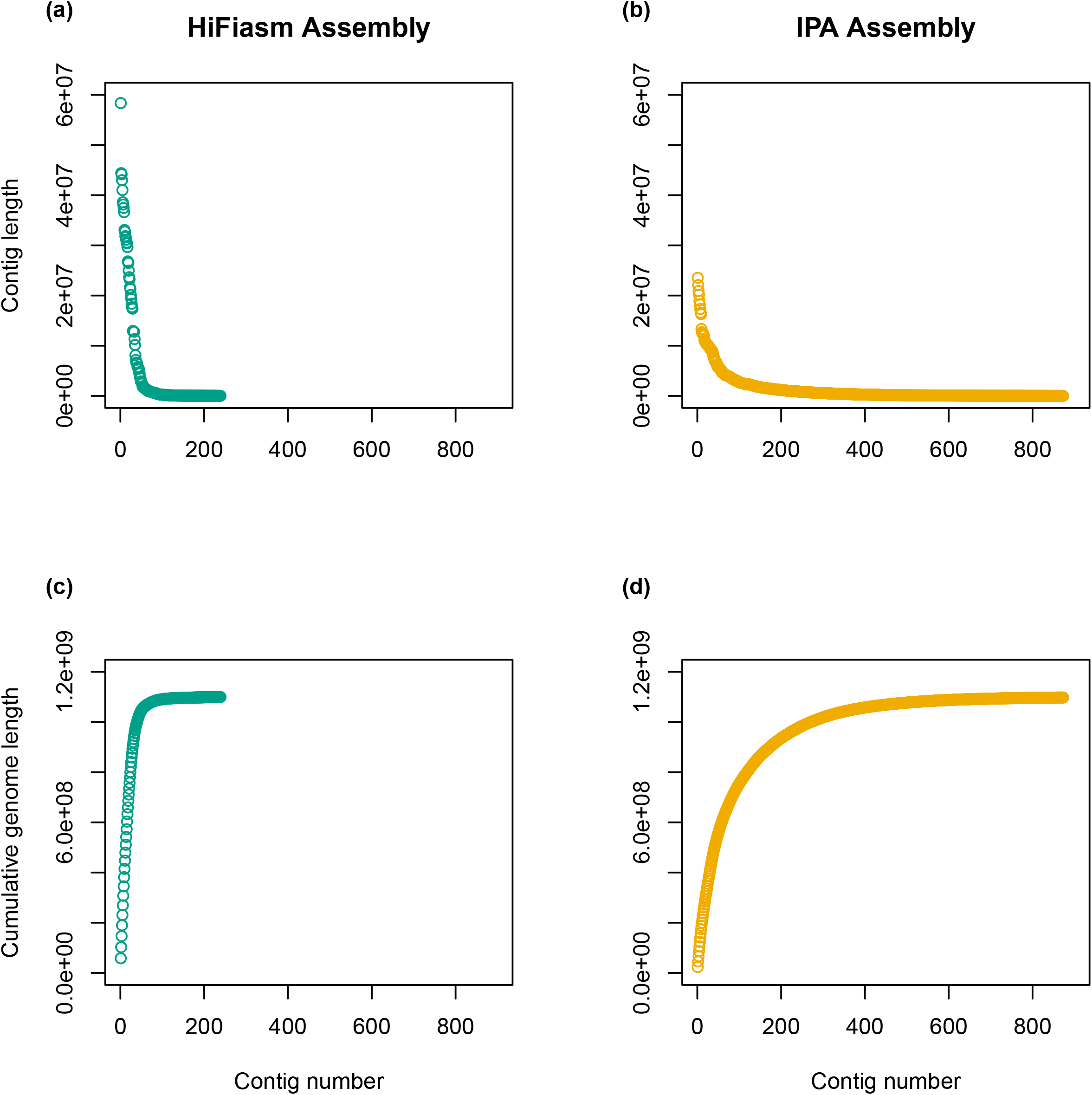
: Visual comparison of the number of contigs, contig lengths, and cumulative genome lengths for both the HiFiasm and IPA genome assemblies. (a) Length of each contig in the HiFiasm assembly. (b) Length of each contig in the IPA assembly. (c) Cumulative genome length of HiFiasm assembly. (d) Cumulative genome length of IPA assembly.

We examined synteny between creek chub and zebrafish, using SynMap, from the platform CoGe (Comparative Genomics, genomevolution.org, Lyons & Freeling 2008). CoGe DAGChainer outputs were used to create circular plots with circos (Krzywinski *et al*. 2009). We used a hard masked version of the genome, uploaded to CoGe (NCBI Window-Masker (Hard) (v1.0,id65989;genomic). The exact zebrafish organism used was Danio rerio (zebrafish;id43752) and the genome was unmasked (v11, id66058; CDS). This is the most recent zebrafish geneome (GRCz11) created by the Reference Genome Constortium, released May 9, 2017 (ncbi.nlm.nih.gov/assembly/GCA 000002035.4). All default analysis and display options were used in the Legacy Version, with the exception of Syntenic Path Assembly (SPA) being selected, contigs without synteny hidden, diagonals coloured by syntenic block, contigs sorted by name, and minimum chromosome size set to 2,830,400bp, which is the length of the 50th largest contig in the creek chub assembly. This SynMap analysis can be generated at any time at this link: genomevolution.org/r/1oxpo. We additionally created a SynMap between only the 25 largest creek chub contigs and the zebrafish genome, by setting the minimum chromosome length to that of the 25 largest contig in the assembly (20,130,130bp, Fig S2). It can be viewed at this link: genomevolution.org/r/1oxpw

We also used SynMap to assess synteny between creek chub and fathead minnow. The zebrafish genome is more complete than the fathead minnow genome (Martinson *et al*. 2022). However, creek chub are more closely related to fathead minnow than to zebrafish (Fig. 1). The version of the genome used was unmasked (v2,id66042;CDS) of GCA_016745375, recently published by Martinson *et al*. (2022). We used the same analysis and display options as mentioned above for the zebrafish SynMap. This SynMap analysis can be regenerated by following this link: genomevolution.org/r/1oxpx. We also created a SynMap with the 25 largest creek chub contigs and the fathead minnow genome (Fig S3). It can be viewed at this link: genomevolution.org/r/1oxq3. SynMap labels are based on the creek chub and fathead minnow genomes’ contig/scaffold codes from the FASTA headers and many of these codes do not intuitively match the contig/scaffold’s corresponding number. See Tables S1 and S2 for a breakdown of the creek chub and fathead minnow genome’s contig/scaffold codes and corresponding contig/scaffold numbers.

We created a simple phylogeny to display the relationship between the creek chub, zebrafish, fathead minnow, and several other model teleost fish (Fig. 1). We created this phylogeny using the R package fishtree (Chang *et al*. 2019), which pulls phylogenetic data from its pre-assembled online database. We added the fish photo with Adobe Photoshop (v22.0.0).

## Results

PacBio HiFi sequencing on two SMRT cells produced 133GB of raw data in FASTQ format. This corresponded to 4,313,794 raw reads with a mean length of 16,406 base pairs, corresponding to a coverage of 64x. An initial genome assembly was constructed by the University of Delaware sequencing facility’s bioinformatics team using Pacific Biosciences’s IPA pipeline, and resulted in an assembly that consisted of 873 contigs, with mean contig length of 1,257,076, and an N50 of 5,722,762 (Table 1). We then improved upon this initial assembly using the HiFiasm pipeline (Cheng *et al*. 2021), resulting in an assembly with 239 contigs, with a mean contig length of 4,599,676, and an N50 of 30,568,897, which is halfway between the N50 of the zebrafish and fathead minnow genomes (Table 1). BUSCO analysis indicates that in addition to being highly contiguous, this genome is largely complete, with a score of 98.0% and 97.9% respectively for the HiFiasm and IPA assemblies (Table 2). BUSCO values were similar between the two assemblies with the exception of a higher proportion of genes designated as complete and duplicated in the HiFiasm assembly relative to the IPA assembly (2.5% verses 1.6%, respectively), however both values are similar to the fathead minnow and lower than the zebrafish (Table. 2). Kraken2 anaylsis did not find any contigs to be entirely contaminated with non-creek chub DNA. As the HiFiasm assembly was comparable to or improved over the IPA assembly in all respects-high completeness and low contamination, but fewer contigs, higher N50, and larger mean contig size-we used the HiFiasm assembly for all subsequent analyses.

**Table 2:**
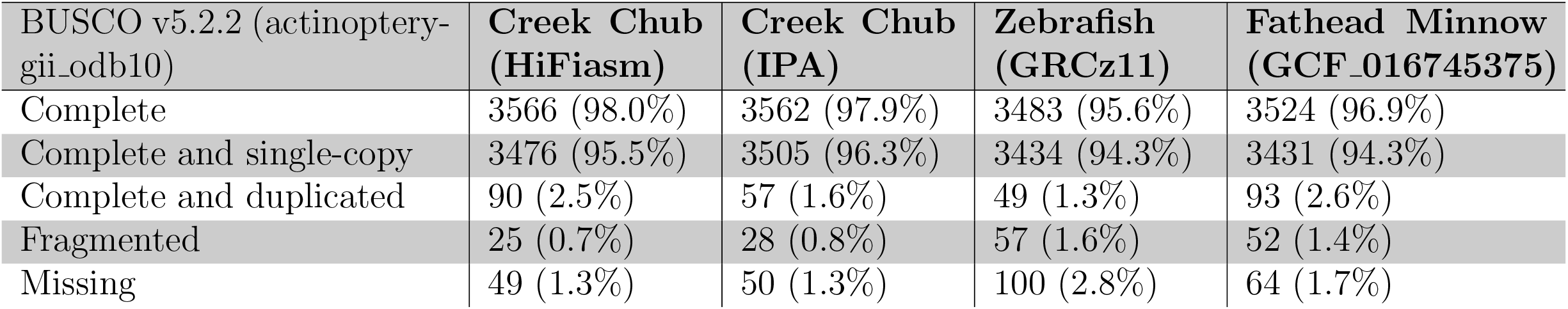
BUSCO (benchmarking universal single-copy orthologs) scores for both the HiFiasm and IPA genome assemblies and the most recent version of the zebrafish and fathead minnow genomes. Generated using BUSCO v5.2.2 (database: actinopterygii odb10). Total number of BUSCO groups searched for each genome: 3640.

Comparative Genomics (CoGe)’s SynMap (Lyons & Freeling 2008) analysis produced 2274 syntenic blocks and 24532 syntenic matches with zebrafish. We see few major chromosomal rearrangements in creek chub relative to zebrafish (Fig. 3). The haploid chromosome number is expected to be the same (n=25) for zebrafish, fathead minnow, and creek chub (Gold & Amemiya 1987), and most contigs in our assembly corresponded in part or whole to zebrafish chromosomes (Fig. 4). While our assembly is less contiguous than the zebrafish genome, in many cases zebrafish chromosomes map to 1–4 larger assembled contigs of the creek chub genome (Fig. 4) and the 50 largest contigs in the creek chub assembly contain just over 95% of the total assembly content (Table 1).

**Figure 3.**
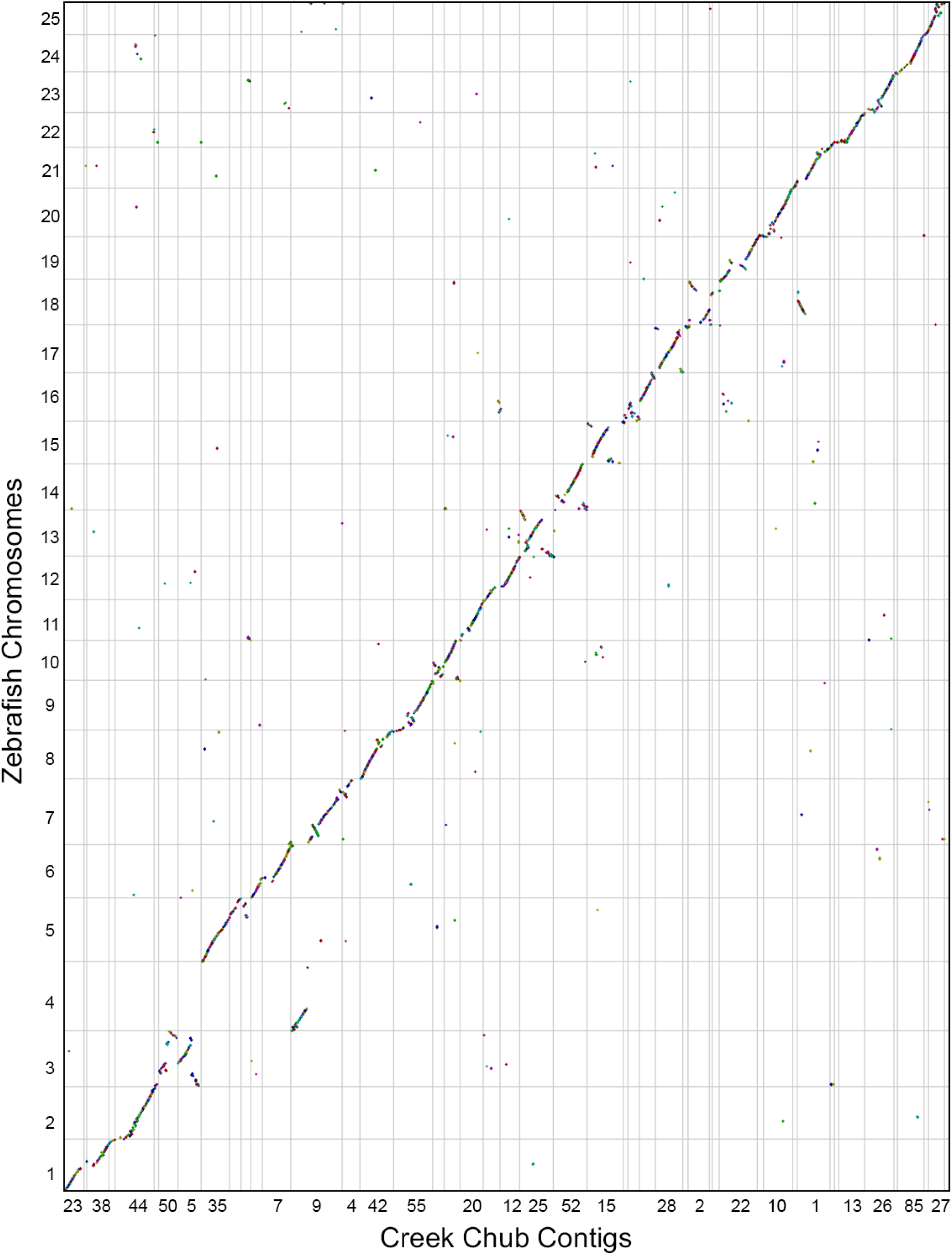
: Dot plot showing synteny between creek chub (x-axis) and zebrafish (y-axis). All 25 zebrafish chromosomes from the GRCz11 version of the genome are present, while only the 50 largest contigs from the creek chub have been displayed, by setting the minimum contig length to 2,830,400 base pairs. The dot plot was made using CoGe’s SynMap (Lyons & Freeling 2008). Each colour represents a different syntenic block. The figure can be regenerated at any time by following this link: genomevolution.org/r/1oxpo

**Figure 4.**
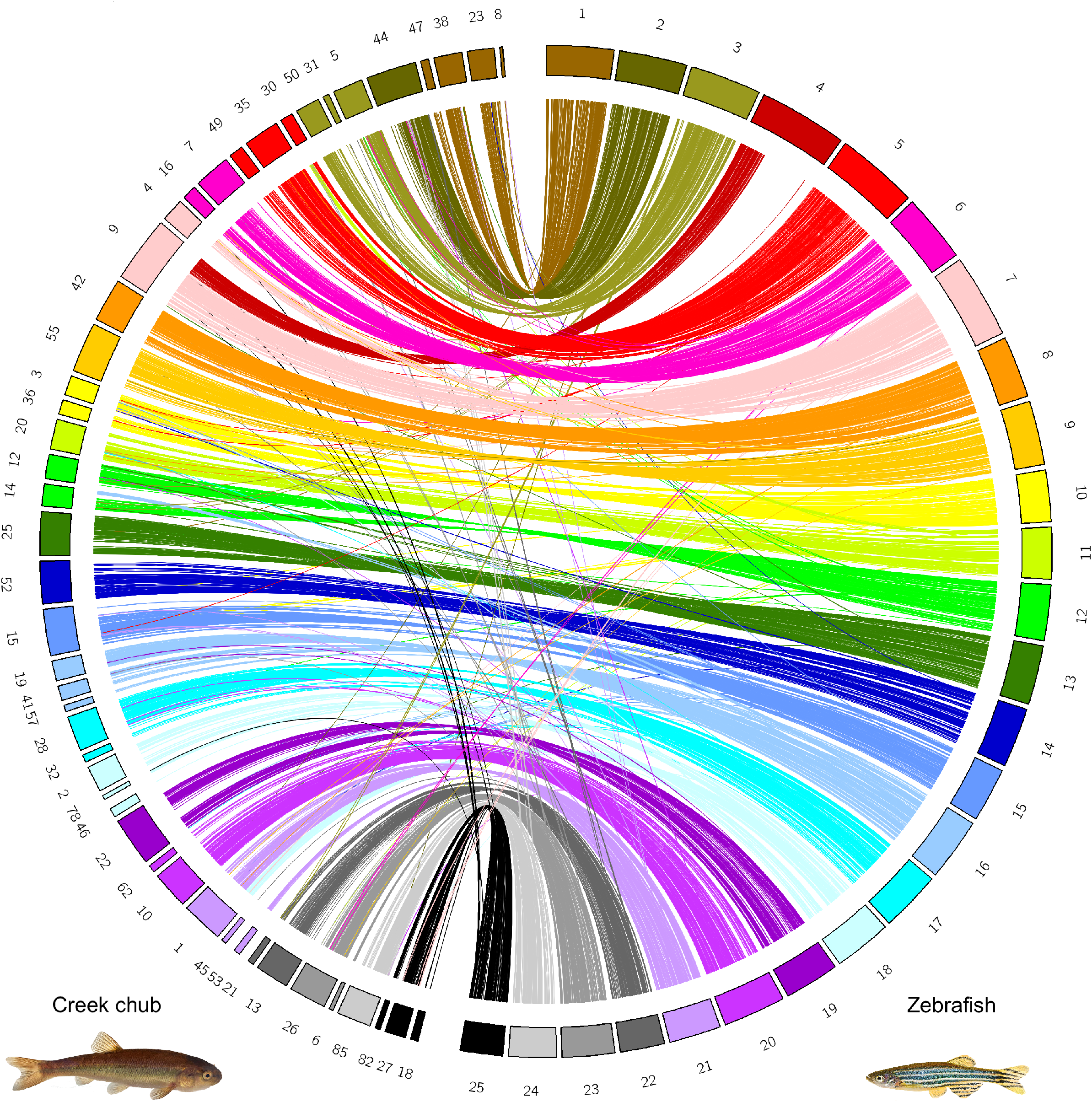
: circos plot of syntenic matches between creek chub (left) and zebrafish (right). Creek chub contigs are coloured and mapped to reflect the zebrafish chromosome it has the majority of synetic matches with. Zebrafish photo credit: Mirko_Rosenau.

Although the large scale pattern is synteny with zebrafish, there are a few regions of the creek chub genome that appear more sharply divergent. In particular, the creek chub contig 9 (the largest complete contig) showed synteny with both chromosomes 4 and 7 in the zebrafish genome (Fig. 4), suggesting there may have been an interchromosomal fusion event in creek chub. Contig 9 also shows two small inversions on the zebrafish chromosome 7 (Fig. 3), indicating intrachromosomal rearrangements. However, as Fig. 3 is coloured by syntenic block, it is obvious that there have been many minor chromosomal rearrangements or mutations. There are no stretches of synteny along any chromosome or contig that are greater than a few dots – with each dot representing a window of 20 genes in which at least 5 genes are syntenic between species. Quantified in a different way, no stretches of synteny along any creek chub contig are greater than 12000 nucleotides, with the average being 407 nucleotides.

In our SynMap analysis between creek chub and fathead minnow (Fig. 5), which produced 19230 syntenic matches in 2042 blocks, we can see that there have also been few major chromosomal rearrangements since these species diverged. One obvious rearrangement is potentially fission or fusion events, whereby scaffold 1 of the fathead minnow genome is split between contigs 28 and 42 of the creek chub genome and scaffold 2 is split between contigs 3, 36, and 44 (Fig. 6). The syntenic matches are less continuous and consistent with some of the fathead minnow scaffolds when compared to zebrafish (Fig. 6 versus Fig. 4). However, this likely reflects the quality of the fathead minnow and creek chub genomes compared to the zebrafish genome, rather than a closer phylogenetic relationship between zebrafish and creek chub than fathead minnow and creek chub. Indeed, creek chub and fathead minnow are more closely related to one another than to zebrafish (Fig. 1). The three species are contained within the order Cypriniformes, with zebrafish in the family Danioninae and fathead minnow and creek chub in the family Leuciscidae (Schönhuth *et al*. 2018, Stout *et al*. 2016). The increased number of syntenic matches over numerous different contigs of each species (Fig. 6), as opposed to contained between a few as we see with zebrafish and creek chub (Fig. 4), are most likely due to the lower continuity and quality of annotation of the fathead minnow genome compared to the zebrafish genome. CoGe predicts syntenic genes based off of sequence similarity, but with a lower quality annotation, is more likely to identify transposable elements or repetitive regions as syntenic between genomes, increasing background noise in the SynMap (Lyons & Freeling 2008).

**Figure 5.**
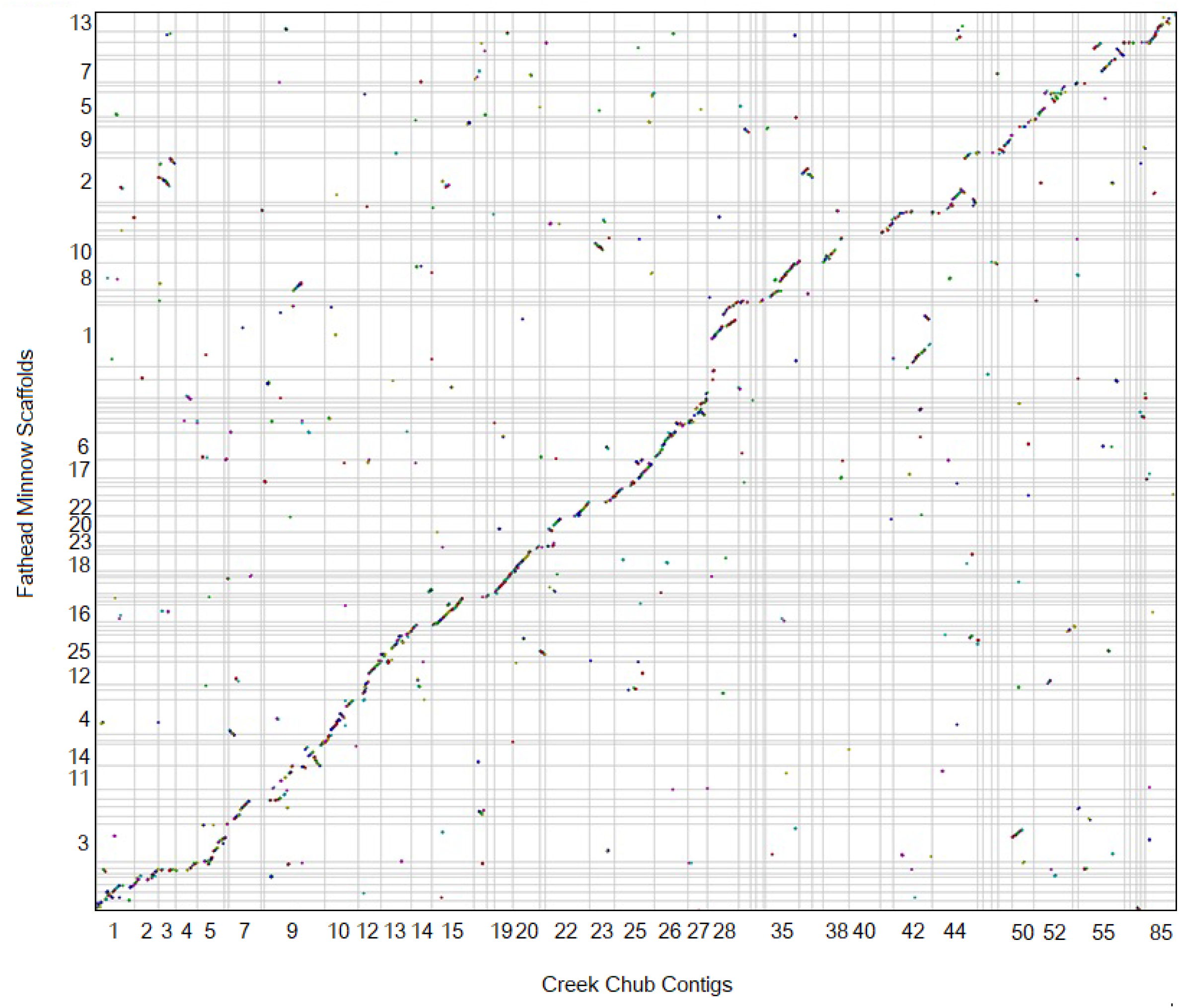
: Dot plot made using CoGe’s SynMap (Lyons & Freeling 2008) showing synteny between creek chub (x-axis) and fathead minnow (y-axis). Only the 50 largest contigs from the creek chub genome have been displayed, by setting the minimum chromosomes length to 2,830,400 base pairs. Each colour represents a different syntenic block. The figure can be regenerated at any time by following this link: genomevolution.org/r/1oxpx

**Figure 6.**
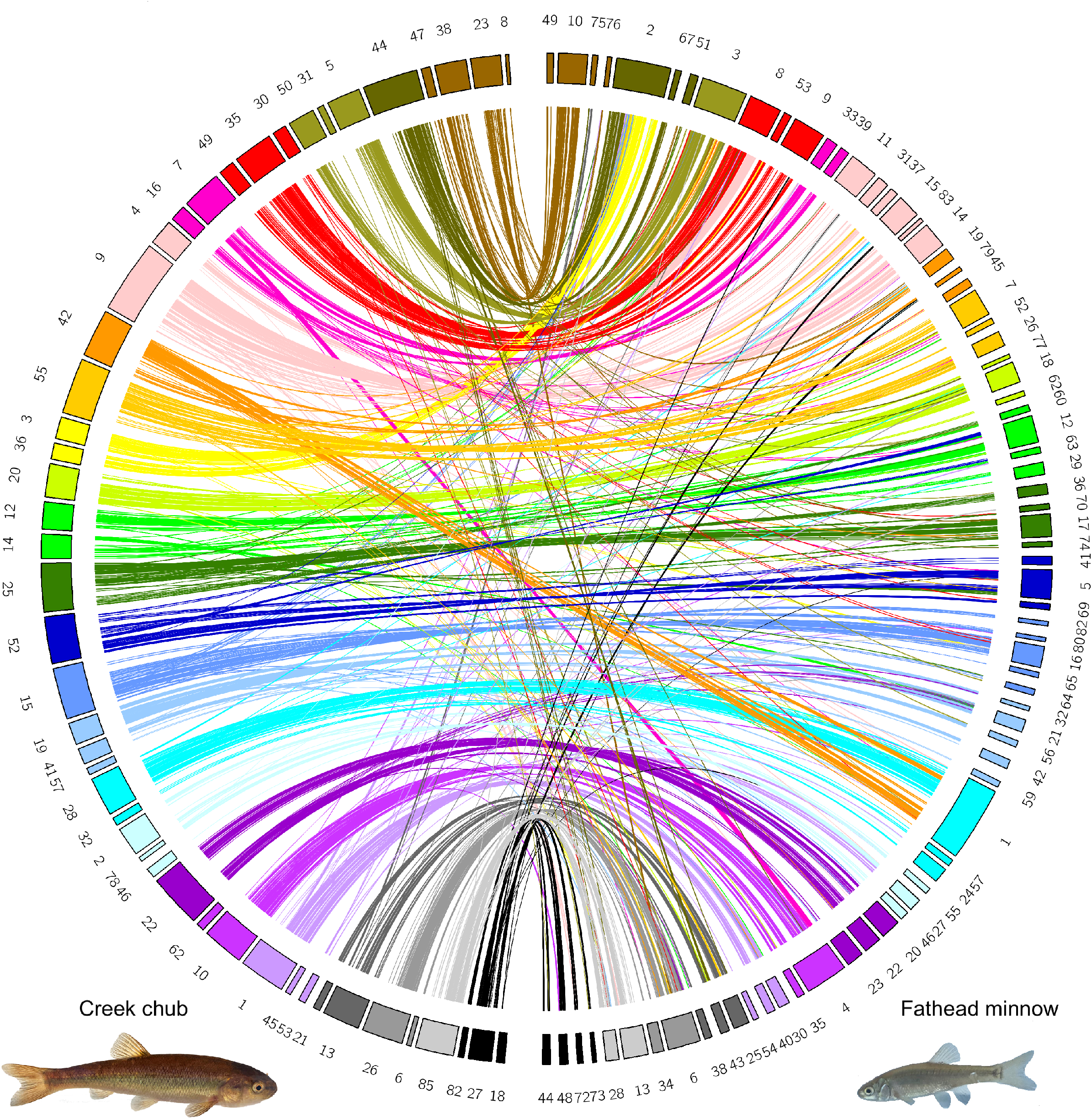
: circos plot of syntenic matches between creek chub (left) and fathead minnow (right). Fathead minnow scaffolds are coloured and mapped to reflect the creek chub contig they have the majority of syntenic matches with.

## Discussion

Our de novo sequencing approach relied entirely on PacBio data, which allowed us to successfully assemble sequence data into a relatively small number of longer contigs (n=239 for the HiFiasm assembly; Table 1). While this assembly is not quite chromosome scale, as the expected haploid chromosome number is 25, there are larger scaffolds which likely approach full chromosomes (Fig. 2), and synteny analyses with zebrafish suggest that each creek chub chromosome is likely covered by 1–4 large contigs (Fig. 4). Analyses of completeness with BUSCO confirm that a high proportion of expected genes are included (about 98% for both assemblies), reinforcing that sequencing produced a high quality reference genome. The contiguity and completeness of this assembly makes it a valuable resource for genomic studies of non-model leuciscid fish. The high contiguity of sequence enabled by PacBio will allow recovery of genetic architecture of traits where linkage of multiple loci might be extremely relevant (for example, examining the genetic basis of sex determination; Meuser *et al*. 2022). A high quality reference genome will also enable analyses that require whole-genome data, such as identifying inversions between closely related species (Faria *et al*. 2019), or demographic inference (MSMC and PSMC; SFS; ABC; Li & Durbin 2011, Schiffels & Durbin 2014, Beichman *et al*. 2018).

In the interest of constructing the most complete and continuous assembly possible from our data, we used two different assembly pipelines, IPA and HiFiasm (Cheng *et al*. 2021). Using HiFiasm, we successfully reduced the number of contigs from 873 to 239, and increased the N50 roughly six-fold (Table 1, Fig. 2). Much of the improvement in N50 and contig number likely resulted from the linking of multiple long contigs to form contigs that approach chromosome length, which enabled better understanding of synteny with other related species (Fig. 4 and Fig. 6).

As expected, much of the creek chub genome is syntenic with previously published genomes of model organisms, namely zebrafish. However, there are also some rearrangements, including a number of inversions and regions that are not syntenic with the zebrafish genome. We do not yet know what functions are encoded by those particular regions of the genome, but structural genome changes, especially of the sex determining region, are likely to play a major role in diversification of species-rich clades of fish like the Cypriniformes (Payseur *et al*. 2018, Huang *et al*. 2020). One particular region to note in the synteny analysis was that approximately half of zebrafish chromosome 4 was not conserved between species. This region is a sex determining region in zebrafish, which has been shown to exist in wild – but not lab raised – strains of zebrafish (Wilson *et al*. 2014). Sex determination systems vary widely across teleost fish species (Bachtrog *et al*. 2014, Pennell *et al*. 2018). Creek chub are not known to have a large sex determining region (Meuser *et al*. 2022) or heteromorphic sex chromosomes (Gold *et al*. 1979), which could be part of the reason why there are no other large regions lacking synteny between the two genomes.

While our creek chub genome assembly does not quite have one large contig per chromosome, for each zebrafish chromosome there are 1–4 larger contigs in our genome assembly that are highly syntenic and likely together comprise the creek chub chromosome (Fig. 4). This tells us that our genome is nearly chromosome-resolution, less a few joins between large contigs. This is especially apparent when comparing to the fathead minnow reference genome; our creek chub reference genome has fewer and larger contigs than the fathead minnow genome (Table 1, Fig. 6). While our creek chub genome is not yet annotated, it is certainly nearly complete and of similar quality to other recently published fish genomes (Martinson *et al*. 2022).

The high-quality creek chub reference genome presented in this paper will enable new insights about the evolutionary history and genome function of leuciscid fish species. Initially, we intend to use this reference genome to investigate the effects of anthropogenic disturbance on a suite of leuciscid fish species. Creek chub and a number of closely related species are widely distributed in North America, and are found in disturbed environments, which makes them an ideal study species for assessing impacts of urbanization and agricultural land use on fish species (similar to previous work in other taxa; Miles *et al*. 2019, Wei *et al*. 2021). A future goal is to produce a genome annotation, which would allow analysis of functional patterns of genomic variation and gene expression in a more meaningful way. More broadly, we are now entering a new and exciting era for genomics of non-model organisms, when it is possible to move beyond using genomes of model organisms as reference, and gain the more fine-grain insights that can only be obtained with a conspecific or closely related reference genome (Gopalakrishnan *et al*. 2017). Generating high quality reference genomes is essential for quantifying genomic variation across the incredible biodiversity of fishes (Fan *et al*. 2020), and will lead to new insights about the evolution of this species-rich group of vertebrates.

## Acknowledgements

Computing was accomplished through an allocation from the Digital Research Alliance of Canada to EGM. We would like to thank T. Frauley for assistance with fieldwork, B. Schultz for productive discussions about analysis of synteny, and S.E. McFarlane for manuscript comments and discussion of what should be in a genome paper. This manuscript was improved by comments from the entire Mandeville lab at University of Guelph. We also thank B. Kingham and O. Shevchenko of the University of Delaware DNA Sequencing & Genotyping Center for coordinating the DNA extraction, library preparation, sequencing, and initial assembly with the IPA pipeline. Finally, we would like to thank to E. Lyons and A. Nelson from CoGe for assistance with creating the SynMap analyses, and the landowners of the property bordering Swan Creek for allowing us access.

## Author contributions

EGM, AVM, and ARP planned the project. ARP and AVM completed field sampling and tissue dissections. AVM and ARP completed the analyses and made the figures, with assistance from EGM. All authors contributed to writing and revising the manuscript.

## Data Availability Statement

Supplemental files are available at FigShare. File S1 contains a photo of the creek chub used to create the reference genome. File S2 contains a syntenic dot plot between zebrafish and the creek chub assembly’s largest 25 contigs. File S3 contains a syntenic dot plot between fathead minnow and the creek chub assembly’s largest 25 contigs. File S4 contains a table of the creek chub assembly’s contig headers, the associated contig number, and length of the contig in base pairs. File S5 contains a table of the fathead minnow assembly’s scaffold code, the associated scaffold number, and length of the scaffold in base pairs. Upon the acceptance of this manuscript, data and scripts used for analysis will be made publicly available in Data Dryad. The genome is available on the NCBI genomes repository, under accession number PRJNA994924. Custom scripts used in this work will be available on Github: github.com/amanda-meuser/CreekChubGenome.

## Conflict of Interest

The authors declare no conflict of interest.

## Funder Information

This research was undertaken using a Resources for Research Groups (RRG) computing allocation from the Digital Research Alliance of Canada. Sequencing for this project was funded by the Canada First Research Excellence Fund, specifically, University of Guelph’s Food From Thought Research Support grant.

## Supplemental Figures

**Figure S1:**
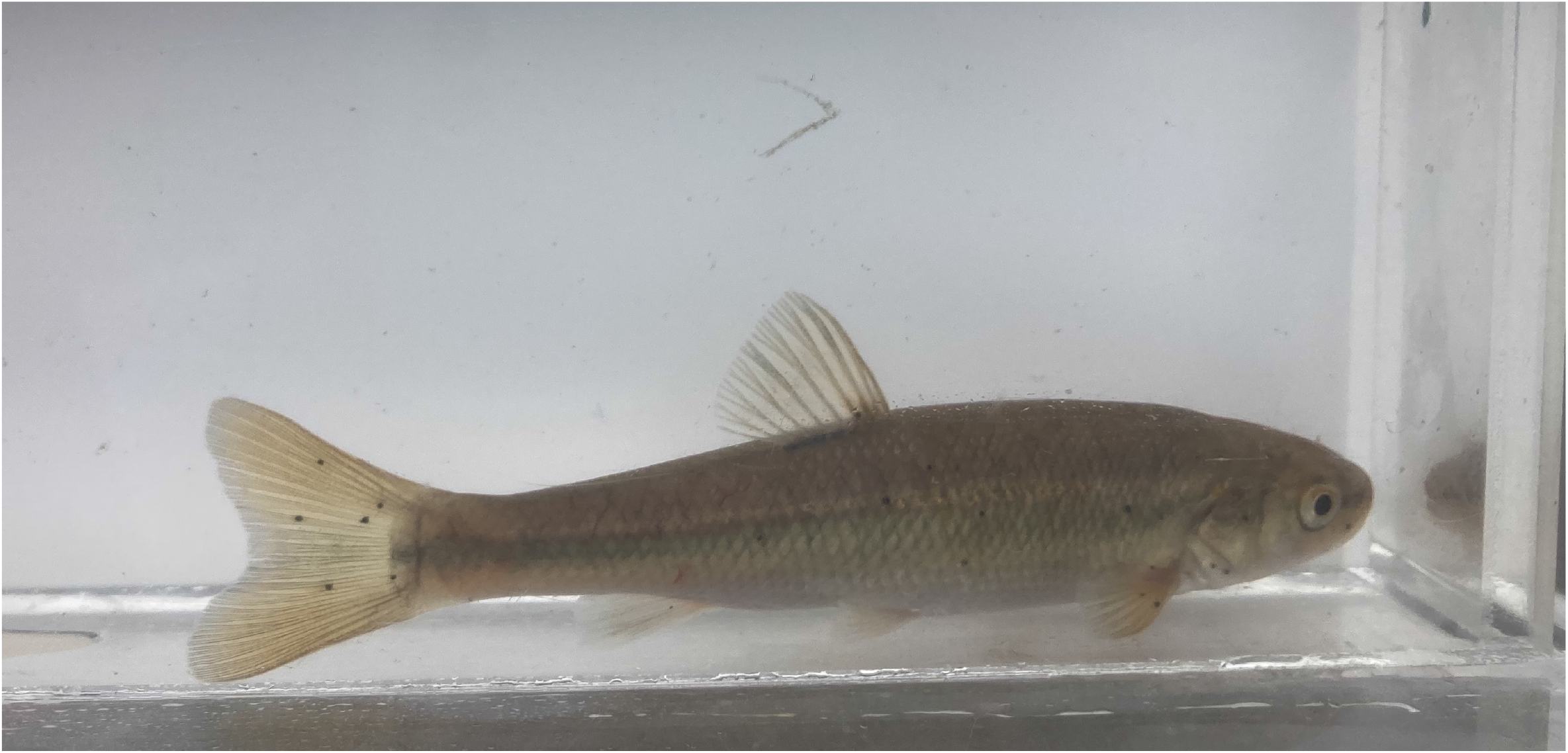
The creek chub individual used to create the reference genome. This fish was sampled from Swan Creek, Ontario, Canada. Note the dot present at the base of the dorsal fin and intermediate scale size compared to similar species. Not visible in the photo are the small barbels in the groove of each side of the mouth and minimally visible is the large mouth and black “moustache”.

**Figure S2:**
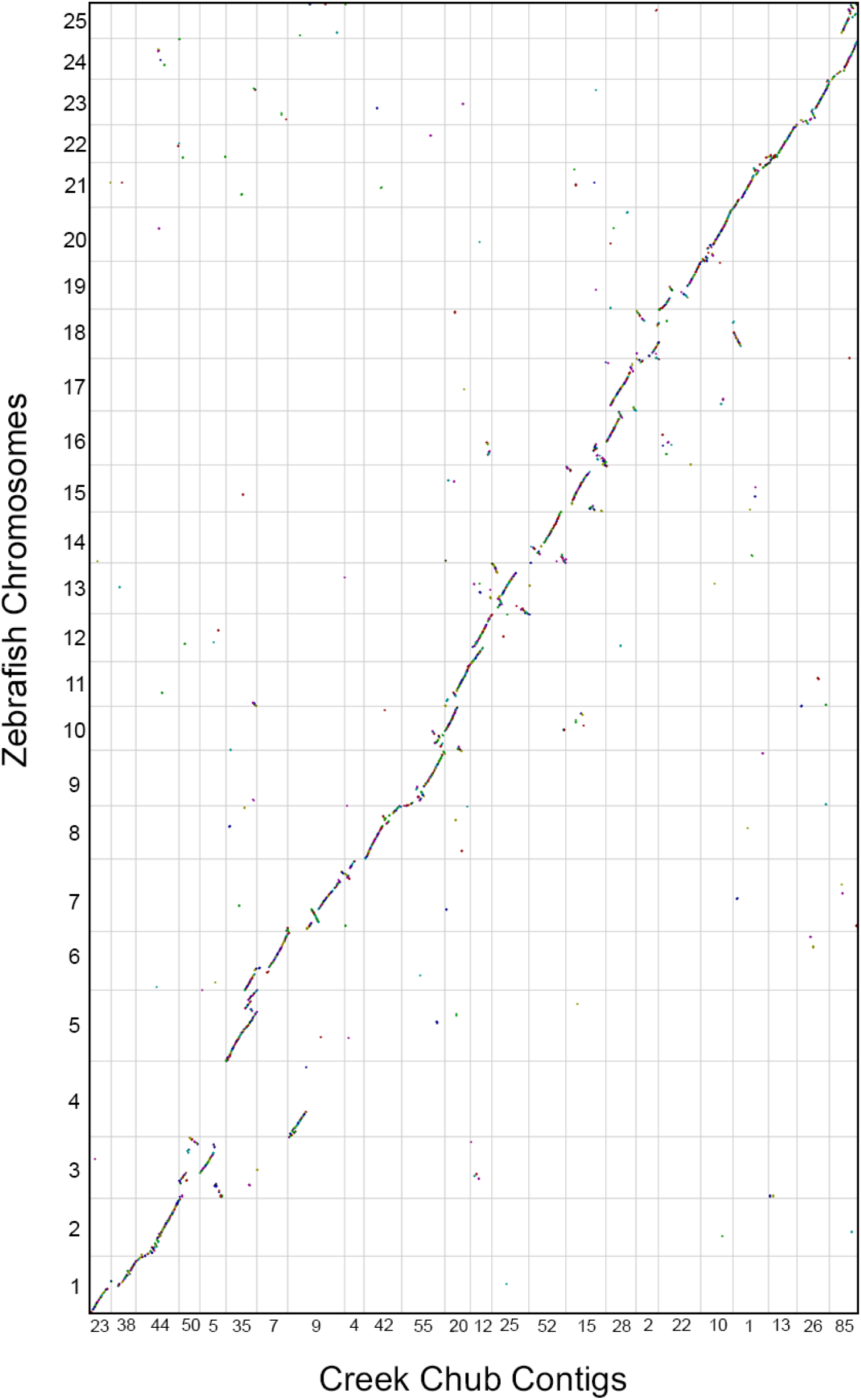
Dot plot made using CoGe’s SynMap (Lyons & Freeling 2008) showing synteny between creek chub (x-axis) and zebrafish (y-axis). All 25 zebrafish chromosomes are present, while only the 25 largest contigs from the creek chub have been displayed, by setting the minimum chromosome length to 20,130,130 base pairs. Each colour represents a different syntenic block. The figure can be regenerated at any time by following this link: genomevolution.org/r/1oxpw

**Figure S3:**
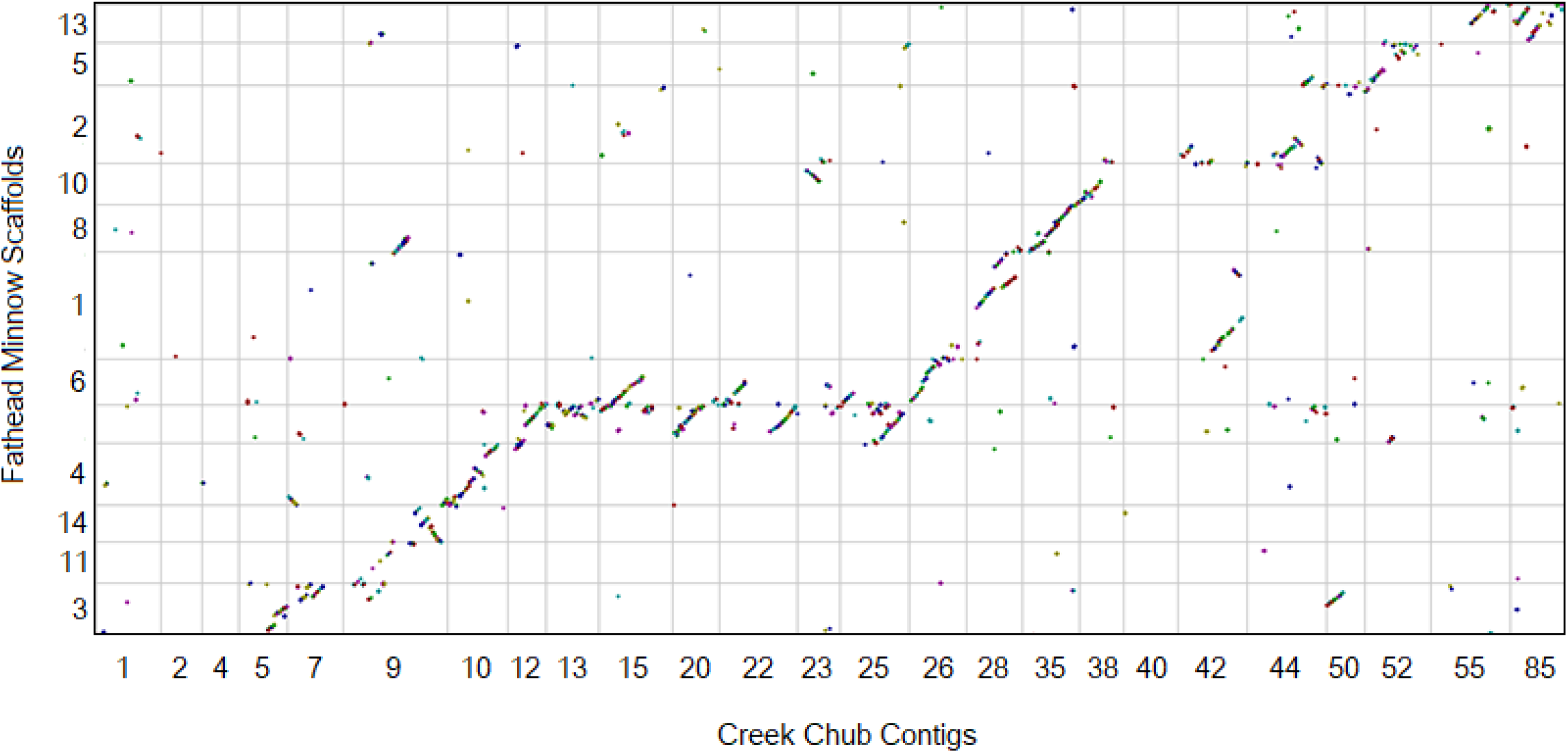
Dot plot made using CoGe’s SynMap (Lyons & Freeling 2008) showing synteny between creek chub (x-axis) and fathead minnow (y-axis). Only the 25 largest contigs from the creek chub genome have been displayed, by setting the minimum contig length to 20,130,130 base pairs. Each colour represents a different syntenic block. The figure can be regenerated at any time by following this link: genomevolution.org/r/1oxq3

## Notes

### Competing Interest Statement

The authors have declared no competing interest.

